# *LEAFY* demonstrates ancestral reproductive functions in the gametophyte and not the sporophyte of the fern *Ceratopteris richardii*

**DOI:** 10.1101/2025.03.21.644678

**Authors:** Hannah McConnell, Jancee R. Lanclos, Katelynn Willis, Nicholas Gjording, Genevieve Stockmann, Julin N. Maloof, Andrew R.G. Plackett, Veronica S. Di Stilio

## Abstract

Flowers are a key reproductive innovation of the angiosperms. They evolved as a modification of the ancestral plant life cycle whereby the haploid gamete-producing generation (gametophyte) became enclosed within the diploid, spore-producing generation (sporophyte). The transcription factor *LEAFY* (*LFY*) initiates angiosperm floral development, yet its lineage predates flowers and is found across all land plants. *LFY* function outside angiosperms is known from the moss *Physcomitrium patens*, where they control the first division of the sporophyte, and from the model fern *Ceratopteris richardii*, a vascular plant without seeds or flowers, where *CrLFY1* and *CrLFY2* maintain vegetative meristem activity. However, how *LFY’*s reproductive role evolved remains unclear. Using over-expression, we uncover new roles for *CrLFY1/2* in fern gametophyte reproduction, particularly in sperm cells and in the gametophyte’s multicellular notch meristem. No sporophytic reproductive function was detected, but over-expression supports a role in fern frond compounding and a conserved role in the zygote’s first division. Our findings highlight an ancestral *LFY* function in fern haploid-stage reproduction, which may have been co-opted into the sporophyte during the origin of the flower.

**Summary Statement:** The origin of *LEAFY*’s floral function is unknown, with only vegetative roles known from seedless plants. We identify ancestral reproductive roles for the first time, unexpectedly in the fern gametophyte.

## Introduction

Flowers are a key innovation of angiosperms, the most recently diverging clade of land plants, that are largely credited with enabling one of the largest evolutionary radiations of all time (Berendse and Scheffer, 2009). The primary role of the transcription factor *LEAFY* (*LFY*) is as a flower meristem identity gene, with loss-of-function mutants producing leaflike structures instead of flowers (Blázquez et al., 1997; Carpenter and Coen, 1990; Molinero-Rosales et al., 1999; Schultz and Haughn, 1991; Souer et al., 1998; Weigel et al., 1992), and constitutive expression resulting in early flowering (Weigel and Nilsson, 1995). *LEAFY* also falls into a special gene class known as pioneer transcription factors, a few of which are known in plants, with roles in developmental reprogramming (Lai et al., 2018; Yamaguchi, 2021). In its pioneer transcription factor role, *LFY* can bind DNA as either a monomer or dimer in heterochromatic regions, activating the expression of downstream genes by directly binding to promoters in heterochromatic regions and through the recruitment of chromatin-remodeling genes (Jin et al., 2021; Winter et al., 2011).

Even though flower meristem identity is considered *LFY’s* canonical role, its homologs are found across land plants, including those without flowers (Sayou et al., 2014), suggesting that its function evolved over time. *LEAFY* can also be active in vegetative meristems, as in certain angiosperms (Kelly et al., 1995; Wang et al., 2008; Zhao et al., 2017; Shu et al., 2000; Moriyama et al., 2024; Souer et al., 1998) and in the shoot apical meristems (SAMs) of gymnosperms (Mellerowicz et al., 1998; Mouradov et al., 1998; Shindo et al., 2001), the angiosperm sister group that produces seeds without flowers. Other functions for *LFY* homologs include regulating axillary meristems in rice (Rao et al., 2008), and compound leaf development in several angiosperms (Busch and Gleissberg, 2003; Champagne et al., 2007; He et al., 2020; Hofer et al., 1997; Jiao et al., 2019; Wang et al., 2013). In the model fern *Ceratopteris richardii* Brongn. (*C. richardii*), expression of a *LFY* paralog was found in developing fronds, and RNAi-mediated knockdown of its two *LFY* paralogs, *CrLFY1/2*, identified roles in maintaining the vegetative meristem of the diploid sporophyte, in embryo development, and the apical cell (stem cell) of early-stage haploid gametophytes. Those results suggest that *LFY*’s vegetative meristem function was present in the most recent common ancestor (MRCA) of ferns and seed plants (Plackett et al., 2018). In the moss *Physcomitrium patens*, a bryophyte, expression of two *PpLFY* paralogs is found in the apical cell of the gametophyte, and their disruption results in the zygote failing to divide into a multicellular embryo, indicating that one or both paralogs are necessary for early sporophyte development (Tanahashi et al., 2005). *LFY*’s involvement in regulating vegetative cell divisions across land plants supports the hypothesis that this function is ancestral, whereas its role in floral meristem identity is derived. However, there is also evidence in support of the role of *LFY* homologs in the reproduction of non-flowering plants. In seedless vascular plants, *LFY* expression has been detected in both vegetative and reproductive organs of the sporophyte (Himi et al., 2001; Rodríguez-Pelayo et al., 2022; Yang et al., 2017), and expression was also seen in reproductive structures (cones) of four conifer genera (Carlsbecker et al., 2013, 2004; Mellerowicz et al., 1998; Mouradov et al., 1998; Vázquez-Lobo et al., 2007). Whether *LFY* also has reproductive functions in *C. richardii* remains unclear, because transgenic knockdown of expression resulted in early termination phenotypes during vegetative development (Plackett et al., 2018). When heterologously expressed in an *Arabidopsis lfy* loss-of-function mutant, a gymnosperm *LFY* ortholog, and the *C. richardii* paralog *CrLFY2* conferred a partial rescue of flower development, where some but not all floral whorls developed, while a moss version did not (Maizel et al., 2005). These experiments suggest that non-angiosperm *LFY* homologs have the potential to perform reproductive functions, motivating further research as the ability to perform functional studies continues to expand beyond seed plants (Di Stilio and Sinha, 2024).

The evolutionary history of *LFY*’s dual vegetative and reproductive meristematic roles presents a compelling question, particularly regarding the emergence of its highly specialized function in floral meristem development. However, few *in planta* functional studies have investigated the potential reproductive role of *LFY* homologs outside of the angiosperms. Ferns are representative of seedless vascular plants, sister to seed plants, and thus an excellent bridge group, with the homosporous fern *C. richardii* representing an effective model (Hickok et al., 1995, 1987; Hickok and Warne, 1998; Plackett et al., 2015; Renzaglia and Warne, 1995). Unlike angiosperms, with their microscopic, short-lived gametophytes, or bryophytes like *P. patens,* where the diploid sporophyte is short-lived, ferns exhibit free-living macroscopic haploid and diploid phases, allowing for investigations into ancestral gene functions in both life stages.

To test whether *LFY* exhibits reproductive activity outside of the seed plants, in either the haploid or diploid stage, we characterized transgenic plants constitutively expressing *CrLFY1* and *CrLFY2* during the development of the fern *C. richardii*, bypassing the issue of developmental arrest at the early gametophyte stage in the RNAi-mediated gene silencing (Plackett et al., 2018). Single and double over-expressors allowed us to investigate the possibility of sub- or neo-functionalization between the paralogs. On the one hand, our findings do not support the hypothesis of *CrLFY* playing a direct role in the reproduction of the sporophyte (sporing), as expected from its angiosperm role. On the other hand, we show new evidence of a reproductive role for *CrLFY* in the gametophyte, via its effect on the notch meristem that produces gametangia, and from the detection of expression in sperm cells. We also generated new functional evidence matching *CrLFY*’s role in the first division of the zygote to that in moss (Tanahashi et al., 2005). Finally, our study further supports *CrLFY*’s role in the regulation of compound leaf development. Taken together, our findings provide the first evidence of a reproductive role for *LFY* in the gametophyte, supporting the hypothesis of evolutionary co-option from this haploid phase to floral meristem identity in sporophytes as the latter became the dominant phase in angiosperms.

## Results

### Fern *LEAFY* misexpression affects frond development in sporophytes

Given that spore production in ferns can be considered analogous to flowering in angiosperms, and that *LFY* overexpression is known to advance flowering, we investigated the hypothesis that *CrLFY* overexpression advances sporing in *C. richardii*. To that end, we generated *35S::CrLFY1*, *35S::CrLFY2* and *35S::CrLFY1+2* transgenic lines (see Materials and methods) and confirmed construct presence (Fig. S1A-B) and overexpression in the leaves (from here on ‘fronds’) of transgenic ferns just before the emergence of sporophylls, the site of meiosis and sporogenesis (Fig. S2A-F, Fig S2 A-B). *35S::CrLFY1* plants had 4.3- to 4.9-fold more expression of *CrLFY1* than wild-type (Fig. S3C), while *35S::CrFLY2* plants had 3.5- to 31-fold more expression of *CrLFY2* than wild-type (Fig. S2E). The time to reproductive transition did not vary between wild-type and transgenic genotypes, i.e., *CrLFY* over-expression did not accelerate sporing (Fig. 1A, *n*=20, *p*=0.72, two-way ANOVA). The number of sporangia produced per square cm and the germination rate of transgenic spores were not affected either (Fig. S3, *n*=6, *p*=0.46, two-way ANOVA). Thus, there was no advancement of reproductive function in the sporophyte of ferns over-expressing *CrLFY*, as predicted from its angiosperm role.

**Figure 1:**
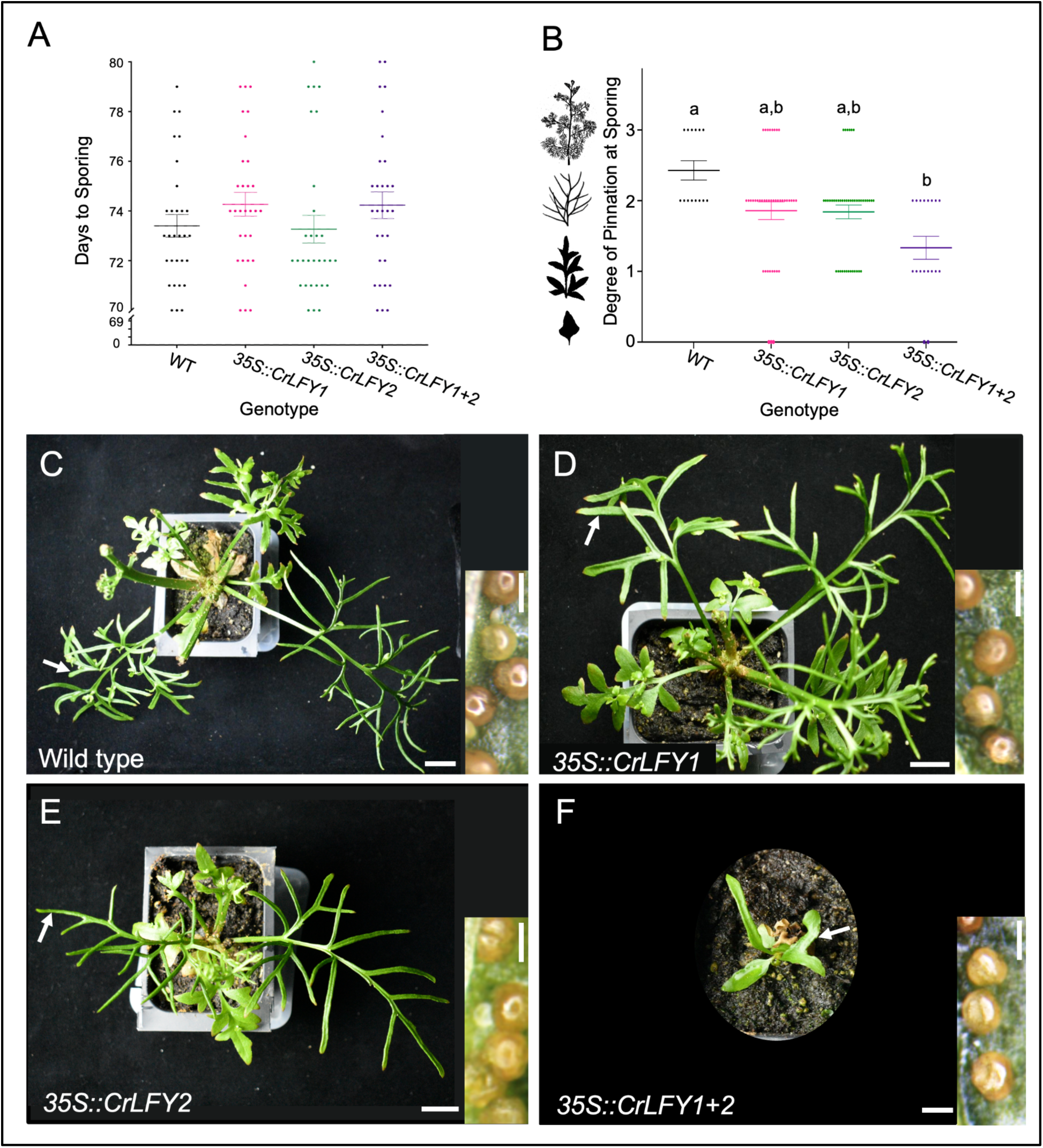
*LFY* orthologs affect frond development but not reproductive transition in *Ceratopteris richardii* sporophytes. Overexpression of *CrLFY1/2* does not change the time to sporing in sporophytes, resulting instead in abnormal development and simpler, less dissected sporophylls (spore-producing fronds) than wild-type at maturity. (A) Days to sporing for wild-type (WT) and transgenic plants (*n*=30, *p*=0.72, two-way ANOVA). (B) The degree of pinnation at sporing for WT and transgenic fronds: 0=simple or lobed, 1=pinnate, 2=bi-pinnate, and 3=tri-pinnate, with representative frond silhouettes shown on the side (not to scale). Different letters denote a statistically significant difference (*n*=30, *p*<0.001, chi-square test). Mean ± standard error of the mean (SEM) shown. Scale Bars = 2 cm. (C-F) Sporophyll pinnation 72 days after fertilization (DAF). (C) WT plant producing sporangia on the abaxial side of bi-and tri-pinnate sporophylls. (D) *35S::CrLFY1* and (E) *35S::CrLFY2* plants producing sporangia on pinnate and bi-pinnate fronds. (F) *35S::CrLFY1+2* plant producing sporangia on pinnate fronds. White arrows mark the sites where sporangia were photographed. Scale bars=1 cm in main panels, or 500 µm in insets.

Wild-type ferns develop compound fronds that become increasingly dissected from simple, lobed, pinnate, bipinnate, to tripinnate (Fig. 1B, frond silhouettes). Reproductive fronds (sporophylls) are bi- or tri-pinnate and produce sporangia, the site of meiosis and sporogenesis (Conway and Di Stilio, 2020, Fig. 1B). *LFY* orthologs have been shown to play a role in *C. richardii* leaf development (Plackett et al., 2018) and leaf compounding in legumes (Hofer et al., 1997). In our experiment, the reproductive fronds, or sporophylls, of *35S::CrLFY1* and *35S::CrLFY2* plants ranged from simple to tri-pinnate (Fig. 1B, D-E), with no statistically significant difference from wild-type ratios (Fig. 1B, C, *n*=20-50, *p*=0.58, chi-square). In contrast, sporophytes overexpressing both *CrLFY* paralogs had sporophylls that were pinnate or bi-pinnate, but never tri-pinnate, and this was statistically significantly different from wild-type expectations (Fig. 1B, F, *n*=30, *p*<0.001, chi-square). Thus, fronds from double over-expressors displayed abnormal development, rather than the expected increased level of dissection from the described role of *LFY* orthologs in compound leaves of certain angiosperms.

### Misexpression of a fern *LEAFY* ortholog alters meristem size in the haploid gametophyte

Given the established role of *CrLFY* in the apical cell (a unicellular meristem) of the fern haploid stage (gametophyte), we tested the hypothesis that it may also be involved in regulating the multicellular “notch” meristem of hermaphroditic gametophytes (whereas male gametophytes are lacking multicellular meristems (Banks, 1997). At 13 days post-sowing (dps) notch meristems of *35S::CrLFY2* and *35S::CrLFY1+2* sexually mature gametophytes both contained, on average, 5 additional meristematic cells compared to wild-type (Fig. 2A-F, *n*=10, *p*<0.001). In contrast, notch meristems of *35S::CrLFY1* gametophytes were not significantly different from wild-type (Fig. 2A-F, *n*=10, *p*=0.51). Additionally, *35S::CrLFY1+2* gametophytes had on average 50.7% larger thalli compared to wild-type, or gametophytes of the same age overexpressing either single *CrLFY1/2* paralog (Fig. 2G, *n*=10, *p*<0.001). Because archegonia (egg-bearing organs) arise from the notch meristem, we quantified these and antheridia (sperm-bearing organs, which arise from mother cells at the periphery of the notch meristem (Banks, 1999) in mature hermaphrodites at 15 dps and 20 dps respectively, when notch-dependent processes could be expected to be even more affected. The average number of archegonia did not vary between genotypes either (Fig. 2I, *n*=30, *p*=0.13, two-way ANOVA) but interestingly, a small proportion of *35S::CrLFY2* gametophytes (3/40 and 1/40, from two independent transgenic lines) failed to develop any archegonia, which was never observed in WT (0/80). Together, these results represent the first evidence for a role of *CrLFY* in the multicellular notch meristem of *C. richardii* gametophytes, while also suggesting functional differentiation between the two paralogs. Wild-type and transgenic plants did not differ in the number of antheridia (Fig. 2H, *n*=20, *p*=0.56, one-way ANOVA), and all gametophytes produced more antheridia in the absence of fertilization, regardless of genotype (Fig. 2H).

**Figure 2:**
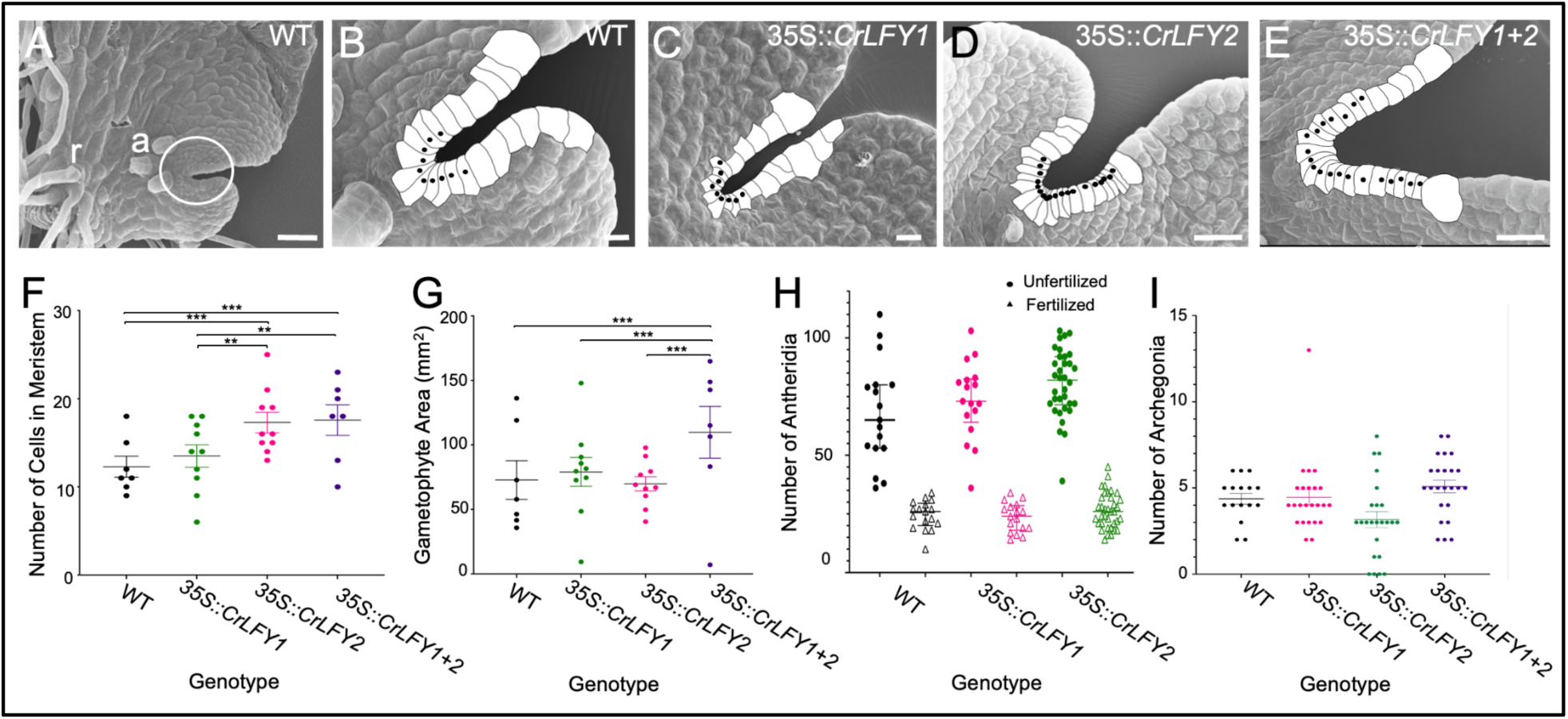
Misexpression of a *LEAFY* ortholog alters the multicellular meristem of the fern haploid stage. *Ceratopteris richardii* gametophytes overexpressing *CrLFY2,* or both paralogs, produce more meristematic cells and are bigger, but with normal amounts of gametangia (antheridia and archegonia). (A-E) Scanning electron microscopy images before maturity (stage Gh6, Conway and Di Stilio, 2020), with notch area cells traced and filled in white, and black dots marking the presumed meristematic cells. (A) WT gametophyte, whole thallus (body) showing the notch area, circled and magnified in (B), three archegonia (a), and rhizoids (r). Notch meristem of representative transgenic gametophytes: (C) 35S::*CrLFY1*; (D) 35S::*CrLFY2*; and (E) 35S::*CrLFY1+2*. Scale bars = 20 µm (A-C), 50 µm (D-E). (F) The number of meristematic cells in WT and transgenic gametophytes at 13 days post-sowing (dps, *n*=10, *** = *p*<0.001, two-way ANOVA). (G) Gametophyte surface area in square mm (*n*=10, *** = *p*<0.001, two-way ANOVA). (H) Number of antheridia at 20 dps in gametophytes that were either flooded at 15 dps or kept from fertilizing (*n*=20, *p*=0.90, one-way ANOVA). (I) Number of archegonia in mature (15 dps) gametophytes (*n*=30, *p*=0.32, two-way ANOVA). Mean ± standard error of the mean (SEM) shown.

### *CrLFYs* are expressed in fern sperm cells

*CrLFY1/2* expression had been previously detected in pooled gametophytes without distinction between the sexes (Plackett et al., 2018). Here, we assessed the sex-specific expression of *CrLFY* in male and hermaphroditic gametophytes before and after sexual maturity. Both paralogs were significantly upregulated in sexually mature males at 15 dps (Fig. 3E), compared to immature males starting to undergo spermatogenesis at 8 dps (Fig. 3C), by 3-fold for *CrLFY1* and by 7-fold for *CrFLY2* (Fig. 3A-B, *n*=3, *p*<0.001, two-way ANOVA). Mature males were also significantly upregulated for *CrLFY* compared to immature hermaphrodites (Fig. 3D), by 6.8-fold for *CrLFY1* and by 7-fold for *CrLFY2*, and to mature hermaphrodites (Fig. 3F), by 12-fold for *CrLFY1* and by 27-fold for *CrLFY2* (Fig. 3A-B, *n*=3, *p*<0.001, two-way ANOVA). Hermaphrodites produce both archegonia and antheridia while males produce only antheridia, in high numbers. Thus, increased expression coincides with the increased number of antheridia containing fully developed sperm in mature male gametophytes. For *CrLFY1*, this result was supported by GUS localization specifically inside the sperm cells using a *CrLFY1_pro_::GUS* transgenic reporter line (Plackett et al., 2018) in both hermaphrodite (Fig. 3E) and male gametophytes (Fig. 3F). Compared to wild-type controls (Fig. 3I-K), GUS staining was found in antheridia undergoing spermatogenesis (Fig. 31), and in mature sperm (Fig. 3J), not in antheridia after sperm release (Fig 3K). Compared to wild-type controls (Fig. 3O, two archegonia shown), GUS activity localized briefly to initiating archegonia (Fig. 3P, archegonium 1) and was not detectable throughout subsequent stages of archegonium development (Fig. 3P, archegonia 2 and 3). To the best of our knowledge, this is the first report of expression of a *LFY* ortholog in sperm, hinting at a potentially novel role in spermatogenesis and/or fertilization.

**Figure 3:**
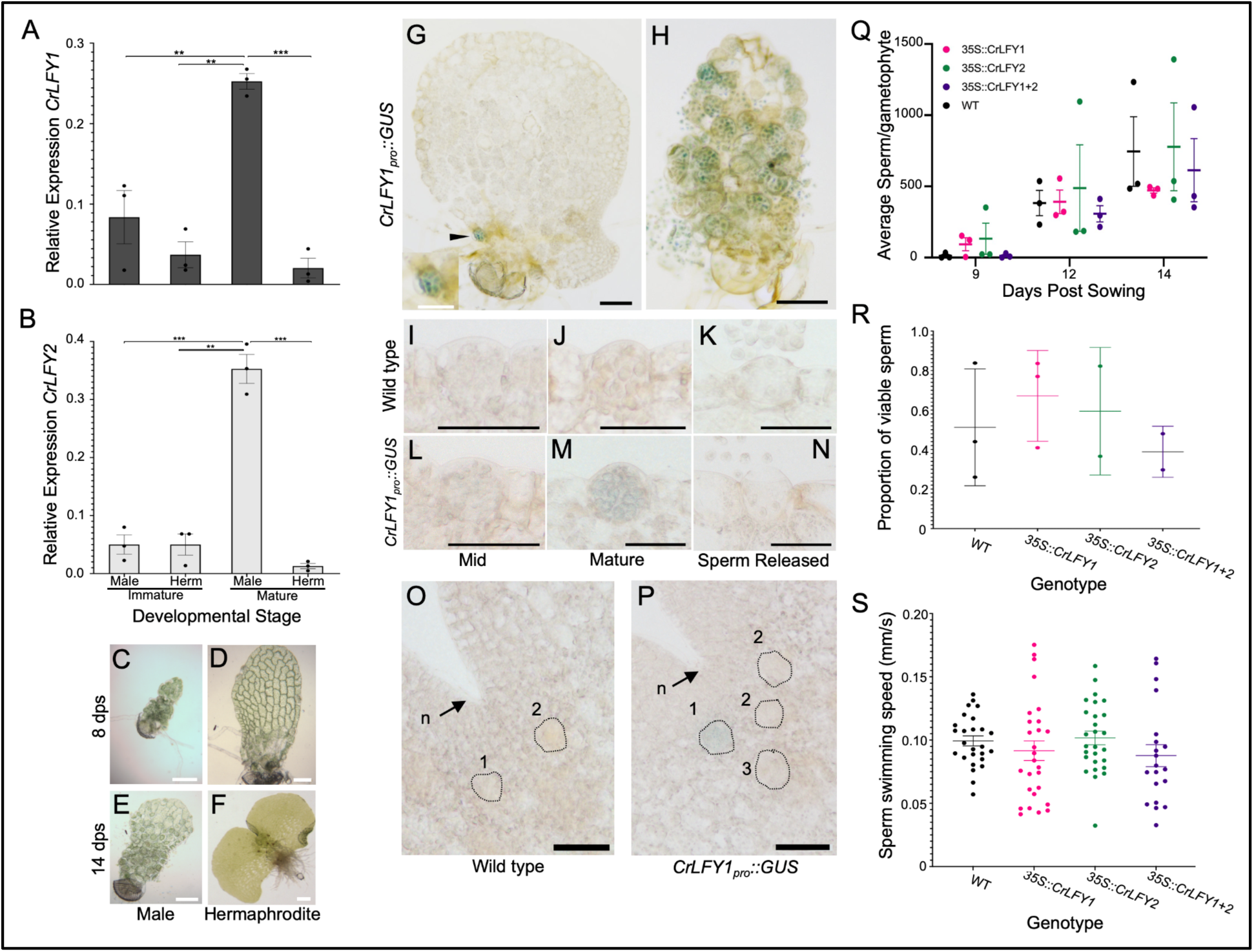
Fern *LFY* paralogs show high expression in mature sperm without a noticeable effect on sperm function when overexpressed. (A-B) Expression of two *C. richardii LFY* paralogs by qPCR (relative to the housekeeping genes *CrACT1* and *CrTBPb*) in gametophytes by sex (male and hermaphrodite, herm) and developmental stage (sexually immature and just starting to undergo spermatogenesis at 8 dps, mature at 15 dps). (A) *CrLFY1* and (B) *CrLFY2* (*n*=3, **=*p*<0.01, *** = *p*<0.001, two-way ANOVA). Representative gametophyte images for the developmental stages used in qPCR: (C) Immature wild-type (WT) male gametophyte at 8 dps, with few immature antheridia. (D) Immature WT hermaphrodite gametophyte at 8 dps. (E) Mature WT male gametophyte at 15 dps, covered in mature antheridia. (F) Mature WT hermaphrodite gametophyte at 15 dps, with antheridia and archegonia. Scale bar=100 µm. (G, H) *CrLFY1_pro_::GUS* expression (blue) in (G) 9 day-old hermaphrodite and (H) male gametophytes. Scale bar=100 µm, or 50 µm (inset). (I-K) Developing antheridia in WT and (L-N) *CrLFY1_pro_::GUS* transgenic gametophytes undergoing spermatogenesis (I, L), with mature sperm (J, M), and after sperm release (K, N). Scale bar=20 µm (O) Archegonia developmental series from the notch meristem (n) in WT and (P) *CrLFY1_pro_::GUS* gametophytes:(1) initiation, (2) development, and (3) fully mature. Scale bar=50 µm. (Q) The average number of sperm cells released from pools of 100 gametophytes at 9, 12, and 14 dps for WT and transgenic gametophytes (*n*=3, *p*=0.6432, two-way ANOVA). (R) The proportion of viable sperm (with propidium iodide as a viability stain) in WT and transgenic gametophytes *(n*=3 reps, 470 to 11140 sperm analyzed per sample, *p*=0.51, mixed-effects binomial). (S) The swimming speed of sperm (in mm/s) for WT and transgenic gametophytes (*n*=25, *p*=0.4126, two-way ANOVA). Mean ± standard error of the mean (SEM) shown.

### Overexpression of *CrLFYs* does not affect sperm function

Because *CrLFY* is expressed in *C. richardii* sperm, we evaluated its putative role in sperm development and/or function. We investigated sperm performance parameters in transgenic gametophytes compared to wild type. There was no difference in the total number of sperm cells produced per gametophyte (Fig. 3Q, *n*=3, *p*=0.64, two-way ANOVA), in sperm viability (Fig. 3R, *n*=3, *p*=0.11, two-way ANOVA), or sperm swimming speed (Fig. 3S, *n*=25, *p*=0.41, two-way ANOVA) between any transgenic line and wild-type. Thus, *CrLFY* misexpression does not appear to influence sperm development or function.

### Misexpression of *C. richardii LEAFY* orthologs disrupts embryo development at the zygote stage

To further investigate the potential role of *CrLFY* in sperm development, we set up controlled fertilization assays in wild-type and *35S::CrLFY1*, *35S::CrLFY2,* and *35S::CrLFY1+2* transgenic lines and recorded fertilization success as the emergence of visible embryos. First, hermaphroditic gametophytes were isolated and flooded with sperm of the same genotype, and followed for 14 days post flooding (dpf); by 10 dpf, 80% of wild-type gametophytes contained visible embryos, and by 14 dpf all showed visible signs of embryo development (Fig. 4A).

**Figure 4:**
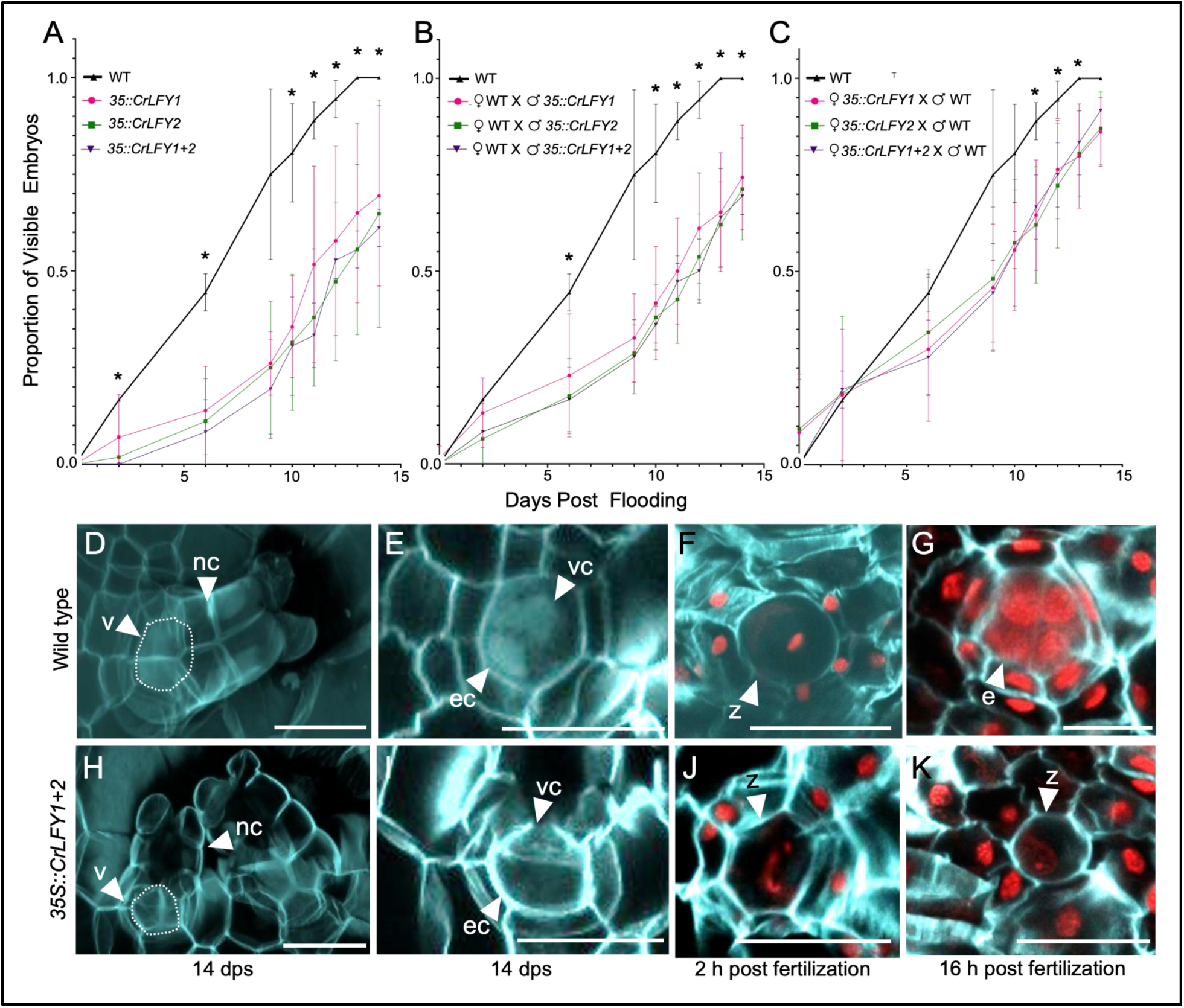
Misexpression of fern LEAFY homologs disrupts early development of the *C. richardii* embryo. In controlled fertilization assays, overexpression of fern *LEAFY* orthologs *CrLFY1/2* prevents zygotes from undergoing cell division and developing into multicellular embryos. Time series of the proportion of gametophytes producing visible sporophytes after flooding (up to 14 dpf) with water containing sperm from the same genotype (A) or outcrossed (B-C). Mean ± standard error of the mean (SEM) shown. (*n*=36, *=*p*<0.05 or less, see Supplemental Table 2 for individual *p*-values). Partial Z-stacks from confocal images comprising the surface of an archegonium in WT (D) and *35S::CrLFY1+2* gametophytes before fertilization (H), stained with SR220 (light blue, cell wall) and Hoechst (light blue, nucleus), denoting the neck canal (nc) and venter (v) cells. White tracing indicates the location of the egg and ventral canal walls, not visible in the partial stack. A single slice taken from the same Z-stack, further zoomed in, denoting the ventral canal (vc) and egg cell (ec, white arrowheads) at the center of the venter, for WT (E) and transgenic gametophytes (I). A zygote (z) is shown at the center of the archegonial venter for WT (F) and transgenic gametophytes (J-K) one day after flooding, stained with SR2200 and Propidium iodide (red nuclei). A WT multicellular embryo (e) developing inside the archegonial venter one day after flooding (G). Scale bar=20 µm.

Transgenic gametophytes for all three constructs had significantly fewer embryos from 10 dpf onwards compared to wild-type, with 36% of *35S::CrLFY1*, 31% for *35S::CrLFY2* and 30% of *35S::CrLFY1+2* gametophytes containing visible embryos 10 dpf (Fig. 4A and Table S2, *n*=36, *p*<0.01, two-way ANOVA). This result suggests that *CrLFY* overexpression disrupts either fertilization or post-fertilization processes.

Second, to determine whether the decrease in visible embryos was due to a maternal or a paternal effect, isolated individual wild-type hermaphroditic gametophytes were flooded with transgenic sperm from each of the three genotypes, and the reciprocal experiment consisted of flooding transgenic gametophytes with wild-type sperm. In crosses where wild-type hermaphrodites were flooded with transgenic sperm, there was a significant reduction in the proportion of gametophytes with embryos after 10 dpf compared to the wild-type x wild-type control (Fig. 4B and Table S2, *n*=36, *p*<0.01, two-way ANOVA), with 46% of *35S::CrLFY1*, 48% of *35S::CrLFY2* and 44% of *35S::CrLFY1+2* hermaphrodite gametophytes containing visible embryos 10 dpf. In contrast, when transgenic gametophytes were flooded with wild-type sperm, there was a significant reduction in the proportion of gametophytes with embryos between 10-13 dpf (*n*=36, *p*<0.01, two-way ANOVA), but by day 14 they experienced a rescue, whereby the proportion of visible embryos was no longer significantly different from wild-type (Fig. 4C and Table S1, *n*=36, *p*=0.06, two-way ANOVA). Additionally, the proportion of transgenic gametophytes fertilized with wild-type sperm containing visible embryos 10 dpf was significantly higher between 10-13 dpf compared to those fertilized with transgenic sperm, whether they were wild-type or transgenic (Supplemental Table 2), with 56% for *35S::CrLFY1* sperm, 57% for *35S::CrLFY2* sperm and 56% for *35S::CrLFY1+2* sperm (*n*=36, *p*<0.05, two-way ANOVA). Thus, the reduced proportion of embryos developing from transgenic fertilizations, along with the partial rescue by wild-type sperm, suggests that *CrLFY* misexpression affects early sporophyte development through both paternal and maternal effects, with a stronger impact when inherited paternally.

To investigate the mechanism underlying the decrease in visible embryos under *CrFLY* misexpression, we examined early embryo development in the archegonia of wild-type and transgenic gametophytes after flooding. Wild-type mature gametophytes produced archegonia with neck cells and a venter containing a ventral canal cell and an egg cell (Fig. 4D-E, Video 1), as expected from prior descriptions (Lopez-Smith and Renzaglia, 2008); *35S::CrLFY1+2* gametophytes showed similar morphology to these controls (Fig. 4H-I). After flooding, fertilized wild-type gametophytes produce a zygote inside the venter, morphologically distinguishable from the egg cell by the increased cell size, taking up the whole ventral canal space (Lopez-Smith and Renzaglia, 2008). Based on these guidelines, we observed that both wild-type and transgenic gametophytes had undergone fertilization to produce a zygote by 2 hrs post flooding (Fig. 4F, J, Video 2). Within 16 hours, 88% of the wild-type gametophytes (22 out of 25) contained a multicellular embryo (Fig. 4G, Video 3) and the rest contained an unfertilized egg cell (Video 4), while only 60% of transgenic gametophytes contained embryos (15 out of 25) and the rest of the transgenic gametophytes exhibited a single cell with an elongated nucleus that we interpret as a zygote arrested before the first cell division (Fig. 4J-K, Video 5). Interestingly, a small proportion of the wild-type gametophytes with multicellular embryos carried a second zygote that had not progressed, suggesting that zygotic arrest may be a normal mechanism to avoid the production of multiple sporophytes per gametophyte (Fig. S4). These results, together with the evidence from our fertilization assays (Fig. 4A-C), suggest that misexpression of either *CrLFY* paralog prevents cell division of the zygote (by an unknown mechanism), causing developmental arrest of the young sporophyte. Thus, this finding suggests a conserved function for *LFY* in early embryo development between the fern *C*. *richardii* and the moss *P. patens*.

## Discussion

As the closest living relatives of seed plants, ferns provide a critical link for understanding *LFY* function. Investigating *C. richardii* bridges the functional gap between angiosperms and bryophytes, offering a framework to trace the evolutionary trajectory of this master regulator in plant development. Here, we asked whether *LEAFY’s* reproductive function in the angiosperm sporophyte (in floral meristem identity, Carpenter and Coen, 1990; Schultz and Haughn, 1991; Weigel et al., 1992) arose from an ancestral vegetative shoot meristem role, or another reproductive function predating the angiosperms. Consistent with prior research indicating that at least one of the *CrLFYs* is necessary to maintain apical meristems in the fern sporophyte (Plackett et al., 2018), our overexpression analysis shows that both *CrLFYs* together regulate frond dissection in the sporophyll. Transgenic plants overexpressing both *CrLFYs* developed less dissected fronds, with fewer pinnae and pinnules (the smallest subdivided segment of a frond), which arise from pinnae initial cells (Hill, 2001), suggesting that *CrLFY* are involved in regulating these apical (stem) cells and that their misexpression inhibits pinna initial identity or activity. Given that *LFY* is found mostly as a single copy gene, and that it can function as both a monomer and a dimer in *Arabidopsis* (Winter et al., 2011), it is unclear at this point whether the compound leaf phenotype is due to an additive effect of increased total *CrLFY* expression or to a regulatory interaction between the two paralogs that would suggest sub- or neo-functionalization. Interestingly, decreased leaf compounding in plants overexpressing *CrLFY* matches that of plants where *CrLFY* was downregulated by RNAi (Plackett et al., 2018), contrary to the predicted increase in dissection based on *CrLFY’s* role in promoting leaf apical cell divisions. This unexpected result suggests that the overall spatial and temporal *CrLFY* expression pattern is more critical than the total expression level in determining pinnae and pinnule outgrowth. Moreover, proper maintenance of pinnae initial identity would require the absence of *CrLFY* expression elsewhere for leaf development to proceed normally. Alternatively, it is also possible that *CrLFY* affects leaf development in a way other than by regulating cell division at pinnae initials. Several angiosperm *LFY* homologs promote compound leaf development, e.g., in legumes (Champagne et al., 2007; He et al., 2020; Hofer et al., 1997; Jiao et al., 2019; Wang et al., 2025); and this role in marginal meristems leading to compound leaves may have arisen separately in ferns and legumes, given the growing consensus that fronds have evolved independently from seed plant leaves (Tsuda, 2024). However, there is also evidence of *LFY* homologs regulating compound leaf development in California poppy, an early-diverging eudicot, (Busch and Gleissberg, 2003), as well as in lotus, an early-diverging angiosperm (Wang et al., 2013), which together with our findings, suggests the alternative hypothesis that this function could represent a deep homology present in the ancestor of ferns and seed plants.

Another unexpected result was the lack of acceleration of the sporing transition in our transgenic plants, given that overexpression of *LFY* accelerates flowering (Weigel et al., 1992). Other fern and lycophyte *LFY* orthologs show increased expression in sporophylls compared to vegetative fronds (Rodríguez-Pelayo et al., 2022). However, such an increase in *CrLFY* expression from vegetative fronds to sporophylls is not found in *C. richardii* (Plackett et al., 2018), and consistent with that, *CrLFY* overexpression did not alter the reproductive characters of sporophylls beyond the described effect on frond dissection. Thus, our results suggest that *LFY’s* floral function did not arise from an ancestral role in the fern sporophyte shoot. Alternatively, although perhaps less likely, a sporing transition role for *LFY* could have been lost in the *Ceratopteris* lineage, or gained in connection with the evolution of heterospory, since *C. richardii* is a homosporous fern and the only available *in situ* hybridization experiments showing expression specifically in sporangia are from heterosporous lycophytes (Rodríguez-Pelayo et al., 2022; Yang et al., 2017).

Our findings also support the capacity of *CrLFY2* to regulate the multicellular ‘notch’ meristem of hermaphrodite gametophytes. In addition to its floral meristem identity role, *LFY* orthologs are also involved in the maintenance of shoot apical meristems in several dicot and monocot angiosperms (Kelly et al., 1995; Moriyama et al., 2024; Shu et al., 2000; Souer et al., 1998; Wang et al., 2008; Zhao et al., 2017), of axillary meristems in rice (Rao et al., 2008), and of marginal meristems in compound leaf development (already described). In gymnosperms, *LFY* paralogs are expressed in the vegetative shoot of *Gnetum* and *Pinus radiata* (Mellerowicz et al., 1998; Shindo et al., 2001), and either one or both *CrLFYs* are necessary to maintain the apical cell in the early fern gametophyte (Plackett et al., 2018). However, this meristem function had not been previously shown in the multicellular notch meristem of the haploid gametophyte stage. That only one of the two paralogs shows an effect when overexpressed suggests the possibility of sub- or neo-functionalization in regulating this meristem. Importantly, while our overexpression experiments suggest that *CrLFY2* contributes to maintaining the notch meristem, we currently lack expression evidence for either paralog at the notch. One possibility is that a similar genetic network regulates apical activity in both the notch meristem and the sporophyte shoot apex, so that overexpression of *CrLFY2* would cause this network to respond, even if the gene is not normally found in the notch. Although the notch meristem is typically considered a vegetative meristem, its association with the development of egg-producing gametangia (archegonia, Banks, 1999; Conway and Di Stilio, 2020; Geng et al., 2022; Hickok et al., 1987), also classifies it as a reproductive meristem, indicating that *CrLFY2* may play a reproductive function in the fern. This hypothesis is supported by the fact that *CrLFY2* has been shown to partially complement an *Arabidopsis lfy* loss-of-function mutant (Maizel et al., 2005), further suggesting that *LFY’s* floral function may have been recruited from an ancestral reproductive role in early, macroscopic gametophytes with multicellular meristems.

In support of the co-option from the gametophyte hypothesis, we detected roles for *LFY* in fern gametophyte reproductive organs. While we did not find differences in the average number of antheridia and archegonia between wild-type and transgenic plants, we observed a small number of *35S::CrLFY2* transgenic plants without archegonia, and we also detected *CrLFY1pro::GUS* signal in early archegonium development. Because the notch meristem triggers the differentiation of archegonia via positional cues (Geng et al., 2022), our data thus suggests that *CrLFY* could influence this differentiation process. *LFY* paralogs are also expressed in the archegonia of the moss *P. patens* (Tanahashi et al., 2005), although loss-of-function mutants did not reveal phenotypes specifically associated with archegonia in that species.

Our findings also identified the novel expression of *CrLFY* in fern sperm, further supporting a reproductive role for *LFY* in the last common ancestor of ferns and seed plants. We determined that in the gametophyte generation, *CrLFY1* and *-2* are most highly expressed in mature male gametophytes, and that in both sexes *CrLFY1* expression is localized to developing sperm within antheridia. *LFY* homolog expression has been reported from antheridia by RNA-seq in the bryophyte *Marchantia polymorpha* (Arnoux-Courseaux and Coudert, 2024; Kawamura et al., 2022), whereas *PpLFY* expression was not detected in moss antheridia (Tanahashi et al., 2005). This is, therefore, the first report of expression of a *LFY* ortholog in male reproductive organs of seedless vascular plants, and the first to show specific sperm cell expression via a *CrLFY1* promoter-driven GUS reporter, suggesting that at least *CrLFY1* may play a role in sperm maturation. Despite evidence of *CrLFY1* expression during spermatogenesis, there was no difference in the number, viability, or swimming speed of sperm cells between wild-type and transgenic lines overexpressing *CrLFYs*, suggesting that whatever role *CrLFY* may play in sperm development, it is not altered by its misexpression. Nevertheless, a proportion (44-48%) of the zygotes derived from eggs fertilized by transgenic sperm were arrested in their development, rather than dividing into multicellular embryos, with a few multicellular embryos present likely from individual differences in the degree of *CrLFY* transgene expression in sperm cells. Overexpression of *CrLFY* therefore appears to prevent cell division of the zygote and arrest its development into a multicellular embryo, and our observations in fertilized wild type suggest that this could be one of the mechanisms by which gametophytes control the number of developing embryos from multiple fertilization events. This phenotype appears to be conserved between bryophytes and ferns, as arrested zygotes were also described in *PpLFY* mutants (Tanahashi et al., 2005), suggesting that an ancestral role in the early development of the sporophyte was retained in the vascular plants, and subsequently lost in the angiosperms (or the seed plants). Even though this role in early development was previously hypothesized based on *CrLFY1pro::GUS* expression in the early embryo (Plackett et al., 2018), the evidence presented here represents the first functional support for *CrLFY’*s role in promoting the development of the zygote. That zygotic arrest occurred in similar proportions across transgenic genotypes, with only 31-35% of zygotes on average developing into embryos at 10 dpf, suggests that both *CrLFYs* are redundant for this role.

The zygotic developmental arrest observed in transgenic plants may be due to alterations to a normal *CrLFY* expression pattern or gradient necessary for the first division of the zygote to progress. The first zygotic cell division in *Arabidopsis*, *Ceratopteris* and *Physcomitrium* is asymmetric (Gooh et al., 2015; Johnson and Renzaglia, 2008; Mansfield and Briarty, 1991; Tanahashi et al., 2005; Ueda et al., 2011; Wang et al., 2020), paralleling animal embryology, where asymmetry is caused by morphogenic gradients within the egg or zygote (Christian, 2012; Jankele et al., 2021). The molecular mechanisms underlying zygote polarity and the first asymmetric division in land plants remain unknown (Matsumoto and Ueda, 2024), but have clearly evolved. In brown algae, sperm entry creates a transient axis (Goodner and Quatrano, 1993), whereas in isolated angiosperm egg cells, the sperm entry point does not affect polarity (Nakajima et al., 2010). Consistent with this observation, *LFY* is not expressed or known to function in the *Arabidopsis* zygote and embryo, and this role has presumably been lost in angiosperms, although *LFY* expression is more restricted in *Arabidopsis* and therefore this aspect deserves further exploration in other angiosperms (Blázquez et al., 1997; Klepikova et al., 2016; Weigel et al., 1992). Native *CrLFY* expression is asymmetric during early fern embryo development (Plackett et al., 2018), but it is unclear whether expression starts at the zygote stage. A possible explanation for the greater severity of zygote developmental arrest in reciprocal crosses with *35S::CrLFY* sperm is that excess exogenous *CrLFY* brought in by the transgenic sperm nucleus during fertilization locally disrupts morphogenic gradients within the zygote after fusion, whereas egg-derived *CrLFY* already present in the cytoplasm is not affected by ectopic expression. Auxin has long been suggested as one such zygotic gradient (Zhang and Laux, 2011), and in *Arabidopsis* floral tissues *LFY* is both regulated by and regulates auxin biosynthesis (Li et al., 2013), providing a potential mechanistic link for the role of *LFY* in seedless land plant embryo development and flowers.

One further possibility involves a role for *CrLFY* in zygotic genome activation (ZGA), the transition of the zygote genome from silent to transcriptionally active (Fu et al., 2024). In *Arabidopsis*, the zygote elongates just before cell division, and at this stage, there is debate as to the degree of contribution from each parental genome (Anderson et al., 2017; Del Toro-De León et al., 2014; Kao and Nodine, 2019; Nodine and Bartel, 2012; Weijers et al., 2001; Xu et al., 2005; Yu et al., 2016; Zhao et al., 2019). Most of the evidence suggests a higher contribution from the maternal genome, however, there is also some degree of paternal contribution to zygotic genome activation (Gehring et al., 2004; Kermicle, 1970; Pignatta et al., 2018). In its function as a pioneer transcription factor in *Arabidopsis*, *LFY* can access regions of heterochromatin that are otherwise inaccessible, working as either a transcription factor and binding to promoters, or recruiting chromatin-remodeling factors to increase activity by other transcription factors (Jin et al., 2021; Yamaguchi, 2021). In mammals, pioneer transcription factors are heavily involved in ZGA (Fu et al., 2024; Kobayashi and Tachibana, 2021), however further investigations will be necessary to determine whether *CrLFYs* also behave as pioneer transcription factors and whether they contribute to ZGA. Although the ultimate mechanism of *CrLFY’s* involvement in zygote division will require further investigation, we propose the testable hypothesis that *CrLFY* expression is tightly controlled in the zygote for cell division to progress, leading to normal embryogenesis, and that any disruptions in this expression pattern, whether via gene silencing or over-expression, will result in developmental arrest.

In conclusion, we show evidence that *LFY* homologs in the fern *C. richardii* regulate meristems in both the haploid gametophyte and diploid sporophyte stages, whereas in seed plants, where gametophytes are ameristic, *LFY* is known to function exclusively in the sporophyte. Moreover, *LFY’s* regulation of zygote development appears to be conserved through at least the non-seed vascular plants. Our findings further support the hypothesis that *LFY’s* derived reproductive meristem role in flowering plants originated in the last common ancestor of ferns and seed plants and was potentially co-opted from the gametophyte to the sporophyte phase as vascular plants became increasingly sporophyte-dominant in their evolutionary history.

## Methods

### Plant materials and growth conditions

All experiments were conducted in wild-type *Ceratopteris richardii* Hn-n accession background (Hickok et al., 1995). Spores were surface-sterilized by a 10-minute treatment of 10% Hypochlorite and 0.1% Tween (Sigma-Aldrich, St. Louis, MO) at room temperature, rinsed four times and then imbibed in sterile MilliQ water for 2-6 days in the dark to synchronize germination (“Dark Start”, Hickok and Warne, 1998). Spores were then sown onto C-fern media at pH 6 in 1% Difco Bacto agar (Carolina Biologicals, Burlington, NC) with 20 µg/mL of Hygromycin B (Millipore Sigma, Burlington, MA) as the selective agent for transgenic plants, and grown in a Percival chamber at 28°C, 16 h light/ 8 hr dark, 80 µmol/m^2^/s fluorescent light under humidity domes. Once gametophytes were sexually mature (Gh7 (Conway and Di Stilio, 2020)), the plates were flooded with 1 mL of sterile water to induce fertilization, and incubated until the resulting sporophytes had 5-6 leaves. Young sporophytes were transplanted to soil (Sunshine #4, Sun Gro® Horticulture, Agawam, MA) in 24-well plug trays kept in standing water in a Conviron Chamber (28°C, 70% humidity, 16 h light/8 h dark) under 80 µmol/m^2^/s fluorescent lights. After approximately 6 weeks, plants were transferred to 10 x 10 cm pots and moved to the greenhouse, kept in standing water, and fertilized every two weeks (Plant Marvel Nutriculture, 20-10-20+, 9.6 g/10 gal). Mature sporophylls were cut and placed in glassine bags to mature for at least three months, after which spores were collected into microcentrifuge tubes and stored at room temperature in the dark.

### Transgenic line preparation

*Generation of Constructs:* The coding sequences (CDS) of *CrLFY1* and *CrLFY2* have previously been validated by cloning (Himi et al., 2001; Plackett et al., 2018). No other *LFY* homologs were detectable by amino acid sequence homology within the *C. richardi*i genome v2.1 (Marchant et al., 2022). Each CDS was amplified from wild-type Hn-n *C. richardii* cDNA by RT-PCR and cloned separately into the *35S::ocs* expression cassette of the pART7 cloning vector (Gleave, 1992) as an *Eco*RI-*Bam*HI restriction fragment. The *35S::CrLFY1::ocs* and *35S::CrLFY2::ocs* constitutive expression cassettes were each cloned separately into the ‘pBOMBER’ *Ceratopteris* transformation vector, carrying a hygromycin resistance cassette driven by the Gateway *nos* promoter (Plackett et al., 2015), as a *Not*I-*Not*I restriction fragment.

*Generation of transgenic lines:* Transformation of 35S::*CrLFY1* and 35S::*CrLFY2* constructs into Hn-n *C. richardii* callus was performed by microparticle bombardment as previously described (Plackett et al., 2015). Each construct was transformed separately. T_0_ sporophyte shoots were regenerated from transformed tissue under 40 !g/ml hygromycin selection. Spores were collected from regenerated plants, and germinated to yield T_1_ gametophytes, which were self-fertilized to produce T_2_ sporophytes after growing on hygromycin and genotyping to confirm construct presence. Double over-expressing transgenic lines were obtained by crossing the two validated T_2_ single over-expressing lines BA14 and BD5, flooding single isolated hermaphrodites overexpressing one paralog with sperm overexpressing the other paralog. The resulting sporophytes were grown on hygromycin plates until transferred to soil and genotyped to confirm the presence of both constructs.

### Validation of transgenic plants

Sporophyte leaves from selfed T_1_ plants were collected at the simple leaf stage (S2, (Conway and Di Stilio, 2020), flash frozen and ground, and genomic DNA was extracted from 10 mg of tissue using a modified CTAB method (Doyle and Doyle, 1987). To validate the presence of the transgene, PCR was performed with construct-specific primers (Supplemental Fig. S1, Supplemental Table S1). Confirmed T_2_ sporophytes were assessed for construct insert number by digital droplet PCR (ddPCR, Supplemental Fig. S2). Primers and probes were designed on the same exon region as the qPCR primers for both genes (Supplemental Table S1). Thermocycling conditions were determined following the BioRad ddPCR Application guide (BioRad, Hercules, CA, USA). After droplet generation and thermocycling, samples were transferred to a QX200 droplet reader (Bio-Rad, Hercules, CA). Droplet counts were analyzed with QuantaSoftTM version 1.7 (BioRad, Hercules, CA) with default or manually adjusted settings for threshold determination to distinguish positive and negative droplets. *CrLFY1* copy number was calculated using *CrLFY2* as the internal reference in each sample, and the reverse was done for *CrLFY2* copy number calculation, with the wild-type reference copy number set to 1 in both cases.

### Phenotyping of transgenic plants

Phenotyping of transgenic plants was carried out in the T2 generation. Isogenic lines were produced by isolating hermaphrodite gametophytes at the G4 developmental stage when hermaphrodites and males can be distinguished (Conway and Di Stilio, 2020), in a 24-well plate and flooding them at stage G7 when they had developed mature gametangia. All transgenic lines were grown alongside WT Hn-n. Phenotype characterization included: spore germination success, quantification of gametangia, gametophyte notch meristem size and cell number throughout development, quantification of archegonia with zygotes/embryos after fertilization, number of pinnae at sporing, pinnae length, days to production of sporangia, and number of sporangia in sporophytes. Statistical analyses included ANOVA, for multiple comparisons, and χ^2^ goodness of fit test for the number of pinnae per sporophyll.

### Gene expression analysis

RNA was extracted from flash-frozen whole single leaves (50-100 mg) or entire young sporophytes using the Spectrum Total Plant RNA kit (Sigma-Aldrich, St. Louis, MO). Primers for *CrLFY1*, *CrLFY2*, and the housekeeping genes *Actin* and *TBP* were as previously designed (Plackett et al., 2018). Primer amplification efficiency was determined with a plasmid serial dilution using the slope of the linear regression line. Primer specificity was tested via melting curve analysis. qRT-PCR of three biological replicates and three technical replicates each were performed in a Bio-Rad CFX Connect with iTaq Universal SYBR Green Supermix (Bio-Rad, Hercules, CA). *CrLFY* expression was calculated using the 2^-ΔΔCt^ method (Livak and Schmittgen, 2001) normalized against the geometric mean of housekeeping gene expression (Hellemans et al., 2007). The standard deviation of the Ct values of each gene was calculated to ensure minimal variation in gene expression. Error bars represent the standard error of the mean for the 2^-ΔΔCt^ values. Relative Expression values of *CrLFY1/2* were compared amongst genotypes by one-way ANOVA followed by Tukey’s comparisons.

### GUS staining

The *CrLFY1_pro_*::GUS transgenic reporter lines used here were previously established (Plackett et al., 2018). Whole gametophytes were stained for GUS as described in (Plackett et al., 2015, 2014), using 1 mg/mL X-GlcA and 20 !M potassium ferricyanide to increase staining strength and specificity. GUS solution was infiltrated into tissue without pre-fixation using a gentle vacuum over two minutes and incubated for 16-24 hours at 37°C in darkness. GUS-stained tissues were cleared by incubating in 70% ethanol solution at room temperature.

### Microscopy

Live gametophytes were photographed on agar using a Q-imaging MicroPublisher 3.3 RTV camera mounted on an Amscope dissecting microscope, or a Zeiss Axio Zoom. For fluorescent microscopy, tissue was fixed in FAA (4% formaldehyde, 5% acetic acid, 50% ethanol) at 4°C overnight. ClearSee (Kurihara et al., 2015) was used for clearing for 7-14 days after fixation. Plants were stained with SCRI Renaissance 2200, SR2200 (Renaissance Chemicals, Selby, North Yorkshire, UK), and either Propidium Iodide or Hoechst 33342 (Thermo Fisher, Waltham, MA). Images were obtained with a Nikon A1R HD25 confocal microscope. Frond and whole-plant photos were taken using a Nikon D3400 hand-held camera with a macro lens attachment. Sporangia quantification was performed by dissecting 2 cm regions of mature sporophylls to reveal sporangia. Images were minimally processed for brightness and contrast using ImageJ (Schindelin et al., 2012). GUS-stained tissue was imaged under Kohler illumination using a Leica DM500 microscope (Leica Microsystems (UK) Ltd., Milton Keynes, UK) mounted with a GXCAM-U3PRO-6.3 digital camera (GT Vision Ltd, Newmarket, UK). GUS staining images were minimally processed for brightness and contrast in Photoshop 2022 (Adobe Inc., San Jose, California, USA).

### Fertilization Assays

WT and *35S::CrLFY* spores were sown on C-fern media as described above. Once gametophytes had developed enough for males and hermaphrodites to be differentiated (stage Gh4, Conway and Di Stilio, 2020), individual hermaphrodites were isolated into 24-well plates, 36 plants per each WT or transgenic line, and the rest of the population was allowed to continue to grow. Once gametophytes were sexually mature (Gh7, (Conway and Di Stilio, 2020), the original plates they had been collected from were flooded with 2 mL water, and 1 mL of post-flood water (containing sperm released from the gametophytes) was collected with a pipette and used to flood the isolated plants. For the first assay, all plants were flooded with water containing sperm of the same genotype. In the second assay, transgenic sperm was used to flood WT gametophytes, and in the third, WT sperm was used to flood transgenic gametophytes. Plants were checked under a dissection microscope for evidence of embryos at 2, 6, and 9 days after flooding, then daily for 14 days. Differences in the proportion of visible embryos were tested by two-way ANOVA followed by Tukey’s comparisons.

### Meristem Quantification

WT and 35S::*CrLFY* gametophytes were fixed in FAA overnight just before sexual maturity (Gh6 (Conway and Di Stilio, 2020)). After overnight fixation in FAA, gametophytes were dehydrated to 100% EtOH through an alcohol series, critical point dried, and sputter coated with gold particles. Samples were imaged on a JEOL JSM-6010 Plus scanning electron microscope and images were analyzed in FIJI v2.10 (NIH, USA) to determine the length and width of cells in the notch meristem area. Hermaphrodite gametophytes transition from dividing from a single apical cell to a multicellular lateral notch meristem as they develop, and because cells that have recently divided periclinally in the notch meristem are narrow and long, we used a 2:1 length-to-width ratio to define meristematic cells. Meristematic cells were counted, and the total surface area of the gametophytes was determined in ImageJ. Differences in the number of meristematic cells, the size of gametophytes and the number of gametangia (antheridia and archegonia) were compared by two-way ANOVA followed by Tukey’s comparisons.

### Sperm performance assays

*Sperm number and viability:* Sperm number was determined from samples acquired by a BD FACSymphony A3 Cell Analyzer (Becton, Dickinson and Company, Franklin Lakes, NJ) using FACSDiva v. 8.0. Sperm viability was determined from samples acquired by a BD Accuri C6 Plus (Becton, Dickinson and Company, Franklin Lakes, NJ). Spermatocytes were stained with propidium iodide and detected with a blue laser of 488 nm. All analyses were conducted in FlowJo v10.9.0 Software (BD Life Sciences) and gated initially on singlet cells. To determine viability, cells were gated to split two distinct propidium iodide staining intensities, where intense staining indicates a non-viable cell.

*Sperm Swimming Speed:* 200 male gametophytes were collected and placed into distilled water. The suspension was pipetted off and placed on a microscope slide with a cover slip. Video of moving sperm was recorded on a Samsung Galaxy A54, mounted on a Leica DM1000 LED compound scope at 10X magnification. Raw videos were processed by DVR-Scan 1.6 software to make a black-and-white mask of the sperm for tracking; these masks were processed by TrackR v0.1.2. Scaled coordinates and frame number (30 frames per second) were used to calculate the speed from frame to frame of the first 10 sperm in view 5 minutes after the initial release of the sperm. These frame-to-frame speeds were then averaged for each of the sperm. Speed differences were compared by two-way ANOVA followed by Tukey’s comparisons.

*Sperm Count:* 100 male gametophytes were collected at differing degrees of maturity – immature (Gm4 (Conway and Di Stilio, 2020)), mature (Gm7 (Conway and Di Stilio, 2020)) and two days after reaching sexual maturity, placed in water and allowed to release sperm for one minute. The suspension was then pipetted off and placed in phosphate-buffered saline. These samples were stored at 4°C for at least 2 weeks. Propidium iodide at a final concentration of 0.4 mg/mL was added 45 min before analysis on the BD FACSymphony (Becton, Dickinson and Company, Franklin Lakes, NJ). Using a HTS plate reader (Bruker, Billerica, MA), samples were analyzed at a flow rate of 1 µl/second. The number of sperm cells was compared by two-way ANOVA followed by Tukey’s comparisons.

*Sperm Viability:* 100 male gametophytes were collected at sexual maturity (Gm7, Conway and Di Stilio, 2020), placed in distilled water, and allowed to release sperm for 60 seconds. The suspension was then pipetted off and placed in MilliQ water. This suspension was filtered through 30 µm nylon mesh (MTC Bio, Sayreville, NJ). Propidium iodide at a final concentration of 0.17 mg/ml was added to the suspension. Spherotech Accucount Blank particles (Biocompare, San Francisco, CA) with a concentration of 10^6^ were added to a final concentration of 77,000. After 6 minutes from gametophytes first being placed in water, samples were analyzed using a BD Accuri C6 Plus (Becton, Dickinson and Company, Franklin Lakes, NJ) at a flow rate of 35 µL per second and a core size of 16 µm. Differences in viability were compared by two-way ANOVA followed by Tukey’s comparisons.

## Acknowledgements

Stephanie Conway and Karen Renzaglia for guidance on *Ceratopteris*-specific microscopy. Jo Bui, Valerie Bentivegna, Maggie Fuqua, and Matt Akamatsu for assistance with ddPCR. Wai Pang Chan and the Biology Microscopy facility (University of Washington), Matt Footer for technical and microscopy support, and Aurelio Silvestroni at the Department of Laboratory Medicine and Pathology Flow Cytometry Core (University of Washington) for technical and experimental design assistance. Anthony Garcia and the Di Stilio lab group for support and feedback.

## Data and resource availability

the datasets in this article are contained in the supplementary materials.

