## Supplementary materials for "*LEAFY* demonstrates ancestral reproductive functions in the gametophyte and not the sporophyte of the fern *Ceratopteris richardii*"

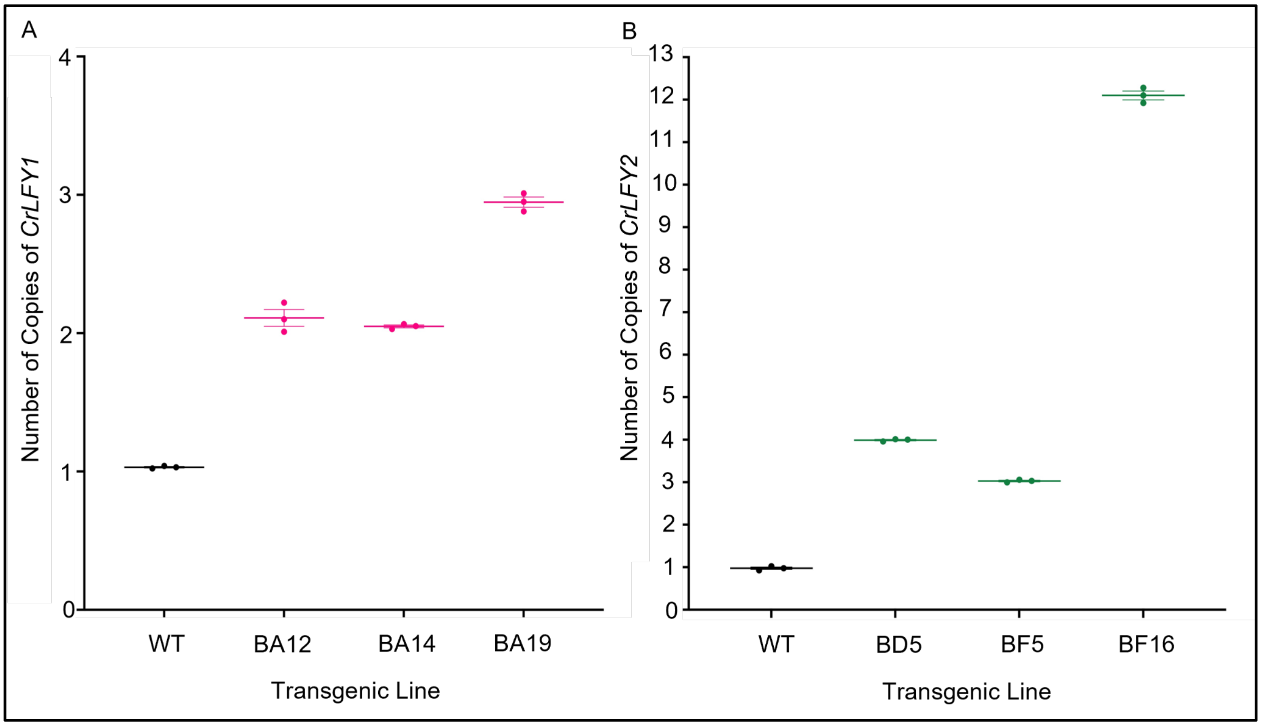


**Figure S1: Insert copy number estimation by digital droplet PCR for the transgenic plant lines characterized in this study.**
(A) Number of *35S::CrLFY1* construct insertions in the three transgenic lines used in this study, compared to wild-type (WT). Two transgenic lines, BA12 and BA14, contain 2 total copies, or one endogenous and one inserted copy of *CrLFY1*, while BA19 contains 3 total copies (i.e., 2 insertions). (B) Number of copies of *35S::CrLFY2* construct insertions in the three transgenic lines used. transgenic line BD5 contains 4 total copies of *CrLFY2*, or one endogenous and 3 inserted, while BF5 contains 3 total copies, and BF16 contains 12. Mean and SEM shown.

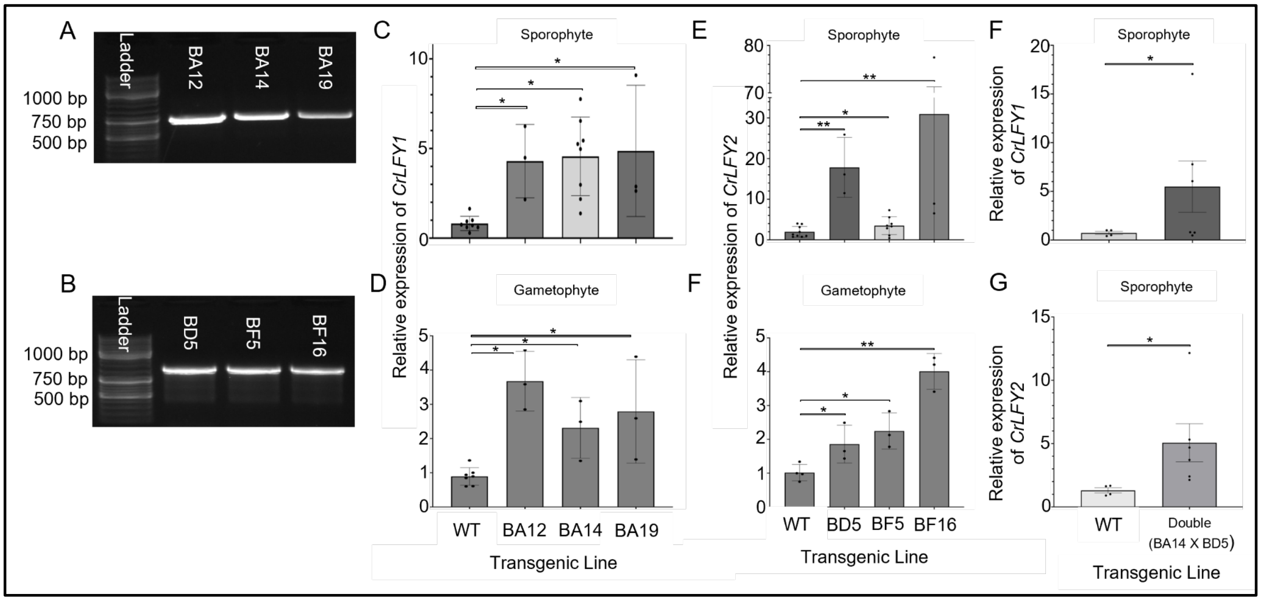


**Figure S2**: **Molecular and expression validation of transgenic plants in this study.** (A) gDNA amplification of three independent transgenic lines per construct characterized here; primers specific for the *35S::CrLFY1* cassette showing bands of the expected size (753 bp). (B) Same for *35S::CrLFY2* transgenic lines (824 bp). (C-D) Relative expression of *CrLFY1* to the housekeeping genes *CrACT1* and *CrTBPb* comparing wild-type (WT) to three independent *35S::CrLFY1* transgenic lines characterized in this study, for (C) sporophytes or (D) gametophytes. (E) Relative expression of *CrLFY2* in WT and *35S::CrLFY2* for (E) sporophytes and (F) gametophytes. *n*=3-8, error bars = standard error of the mean (SEM). Expression of *CrLFY1/2* was significantly higher in transgenic plants compared to wild-type controls (* = *p*<0.05, **=*p*<0.01, one-way ANOVA). See Table S1 for primer sequences.

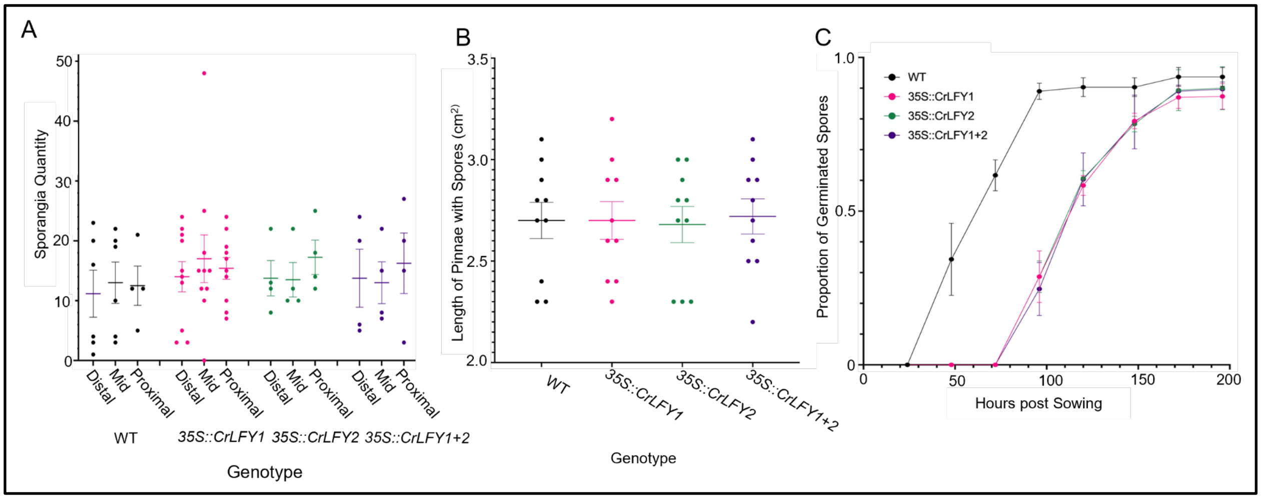


**Figure S3: Number of sporangia, length of sporophyll pinnae, and spore germination success in wild-type (WT) and transgenic *C. richardii*.** (A) The average number of sporangia found across the tip, middle, and bottom regions of a frond ($n$=5-10, $p$=0.4371, two-way ANOVA); (B) Length of individual pinnae from sporophylls bearing sporangia ($n$=10, $p$=0.9917, two-way ANOVA); and (C) Proportion of WT and transgenic spores that germinate 200 hrs after sowing, out of 50 from 3 independent plates ($n$=3, $p$=0.4583, two-way ANOVA). Differences at early time points attributed to a delay in germination due to growth on Hygromycin B-containing media. Mean values ± standard error of the mean (SEM) shown.

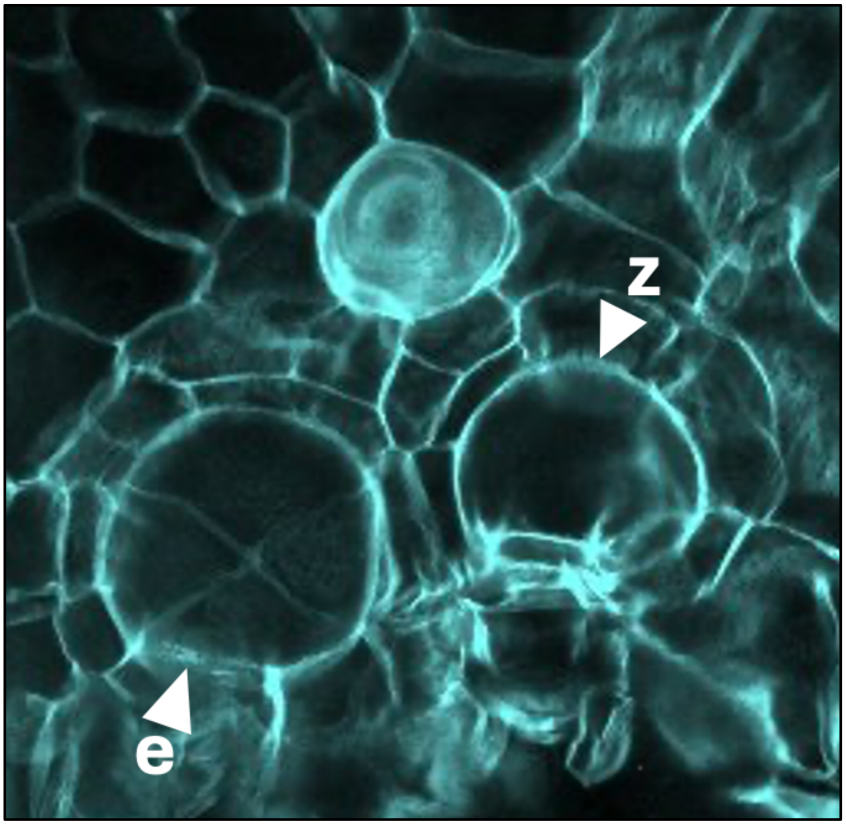


**Figure S4: Example of secondary fertilization in wild-type gametophytes, with multicellualr embryo and an arrested zygote.** Wild-type gametophytes produce a multicellular embryo (e) within 16 h of fertilization. Representative photo from low-frequency secondary fertilization, with a multicellular embryo (4 cells visible, but likely at 8-cell stage), and a single cell zygote (z) that will not develop into a sporophyte beside it.

| **Primer name** | **Sequence 5’→ 3’** | **Purpose** |
|---|---|---|
| qCrLFY1F | ACA AGC ATG CTA TTA TCC ATT GGT | qPCR gene expression |
| qCrLFY1R | TCA CTG TCC TTG CTC TTC TCT AAA | qPCR gene expression |
| qCrLFY2F | GAC TCC TTG CTC TAC CTG AAC CTA | qPCR gene expression |
| qCrLFY2R | CTT CAC CAG GCT CTG TCA CTA TAA | qPCR gene expression |
| qCrACT1_F | GAG AGA GGC TAC TCT TTC ACA ACC | qPCR reference gene |
| qCrACT1_R | AGG AAG TTC GTA ACT CTT CTC CAA | qPCR reference gene |
| qCrTBPb_F | ATG AGC CAG AGC TTT TCC CC | qPCR reference gene |
| qCrTBPb_R | TTC GTC TCT GAC CTT TGC CC | qPCR reference gene |
| HygF2 | CTTCTACACAGCCATCGGTC | Transgenic validation |
| HygR | CCGATGGTTTCTACAAAGATCG | Transgenic validation |
| CrLFY1_F | AAATAGGGCCACCTGGACTC | Transgenic validation |
| CrLFY1_R | CATTCTTCTTTCCCCTTGCC | Transgenic validation |
| CrLFY2_F | TGTAGAAGGCACCAGGGAAC | Transgenic validation |
| CrLFY2_R | TCCCCGTCCTCACCAGGCTC | Transgenic validation |
| 35S_end | AAACCTCCTCGGATTCCATT | Transgenic validation |
| OCSend | TTAGAATGAACCGAAACCGG | Transgenic validation |
| CrLFY1_4,707 F | ACCTGGACTCCTGGCTCTAC | ddPCR insert number |
| CrLFY1_4,874 R | TTTCCCCTTGCCACTTCACC | ddPCR insert number |
| CrLFY2_1,868 F | ACTGCTGCTCAGAATGGTCCC | ddPCR insert number |
| CrLFY2_2,070 R | TCTCTGGTCCTGTCATCCCC | ddPCR insert number |

**Table S1**: Primers used in this study (qPCR primers from Plackett et al., 2018).

| Sperm Contribution | Construct | Day 2 | Day 6 | Day 9 | Day 10 | Day 11 | Day 12 | Day 13 | Day 14 |
|---|---|---|---|---|---|---|---|---|---|
| Same genotype | 35S::*CrLFY1* | **<0.05** | **<0.01** | 0.10 | **<0.05** | **<0.01** | **<0.01** | **<0.01** | **<0.01** |
| | 35S::*CrLFY2* | **<0.01** | **<0.01** | 0.07 | **<0.01** | **<0.001** | **<0.01** | **<0.01** | **<0.05** |
| | 35S::*CrLFY1+2* | **<0.001** | **<0.05** | 0.06 | **<0.05** | **<0.01** | **<0.01** | **<0.01** | **<0.01** |
| Transgenic flooding WT | 35S::*CrLFY1* | 0.39 | **<0.01** | 0.13 | **<0.05** | **<0.001** | **<0.001** | **<0.001** | **<0.001** |
| | 35S::*CrLFY2* | **<0.05** | **<0.01** | 0.11 | **<0.05** | **<0.001** | **<0.001** | **<0.001** | **<0.01** |
| | 35S::*CrLFY1+2* | 0.37 | **<0.05** | 0.09 | **<0.05** | **<0.01** | **<0.01** | **<0.05** | **<0.05** |
| WT flooding transgenic | 35S::*CrLFY1* | 0.98 | 0.070 | 0.24 | 0.09 | **<0.05** | **<0.05** | **<0.01** | 0.07 |
| | 35S::*CrLFY2* | 0.98 | 0.25 | 0.27 | 0.11 | **<0.05** | **<0.05** | **<0.01** | 0.07 |
| | 35S::*CrLFY1+2* | 0.65 | 0.14 | 0.23 | 0.11 | **<0.05** | **<0.05** | **<0.05** | 0.08 |
| WT flooding transgenic vs Same genotype | 35S::*CrLFY1* | 0.43 | 0.17 | **<0.05** | **<0.05** | 0.57 | **<0.05** | **<0.05** | **<0.05** |
| | 35S::*CrLFY2* | 0.24 | 0.03 | 0.12 | **<0.05** | 0.06 | **<0.05** | **<0.05** | **<0.05** |
| | 35S::*CrLFY1+2* | 0.07 | 0.27 | 0.33 | **<0.05** | **<0.05** | **<0.05** | **<0.05** | **<0.05** |

**Table S2**: Statistics (*p*-values) for the proportion of multicellular embryos observed up to two weeks after fertilization (flooding) from two-way ANOVA. The proportion of multicellular embryos for each genotype was compared to the proportion of multicellular embryos found in wildtype on each day, $n$=36, bold indicates a statistically significant difference.
